# A dietary sterol trade off determines lifespan responses to dietary restriction in *Drosophila melanogaster*

**DOI:** 10.1101/2020.08.21.260489

**Authors:** Brooke Zanco, Christen K. Mirth, Carla M. Sgrò, Matthew D.W. Piper

## Abstract

Diet plays a significant role in maintaining lifelong health. In particular, lowering the dietary protein : carbohydrate ratio can improve lifespan. This has been interpreted as a direct effect of these macronutrients on physiology. Using *Drosophila melanogaster*, we show that the role of protein and carbohydrate on lifespan is indirect, acting by altering the partitioning of limiting amounts of dietary sterols between reproduction and lifespan. Shorter lifespans in flies fed on high protein : carbohydrate diets can be rescued by supplementing their food with cholesterol. Not only does this fundamentally alter the way we interpret the mechanisms of lifespan extension by dietary restriction, these data highlight the important principle that life histories can be affected by nutrient-dependent trade-offs that are indirect and independent of the nutrients (often macronutrients) that are the focus of study. This brings us closer to understanding the mechanistic basis of dietary restriction.

## Introduction

Dietary restriction, also called calorie restriction, is a moderate reduction in food intake that extends healthy lifespan across a broad range of taxa, from yeast to primates (Chapman & Partridge, 1996; Colman et al., 2009; Lin et al., 2002; McCay *et al.*, 1935). The generality of this observation has inspired confidence that the health benefits of dietary restriction might also be employed to improve human ageing (Campisi et al., 2019). In an attempt to harness its benefits, a great deal of current research is focused on discovering the nutritional components and the molecular mechanisms that underpin the lifespan benefits of dietary restriction (López-otín *et al.*, 2013; Simpson et al., 2017).

Our current understanding of how diet modifies lifespan has grown out of evolutionary theory and experiments using model organisms. The most prominent theoretical explanation has been the disposable soma theory, which employs resource-based trade-offs to explain how dietary restriction can benefit lifespan (Kirkwood, 1977; Shanley & Kirkwood, 2000). This theory postulates that organisms will maximise fitness by strategically partitioning limiting dietary energy either to reproduction or somatic maintenance, the latter determining lifespan. This means that longer lifespan is inevitably coupled with reduced reproduction because both traits compete for the same limiting resource.

Recent experimental work across a broad range of taxa has challenged the disposable soma theory by demonstrating that reproduction and lifespan respond predominantly to the balance of dietary macronutrients, not the overall energy content of the diet (Mair *et al.*, 2005; Lee *et al.*, 2008; Skorupa *et al.*, 2008; Grandison et al., 2009; Solon-Biet *et al.*, 2014; Solon-Biet *et al.*, 2015; Simpson *et al.*, 2017; Regan *et al.*, 2020). Specifically, high protein, low carbohydrate diets are consistently associated with high reproduction and short lifespan, while low protein, high carbohydrate diets are associated with longer lifespan and lower levels of reproduction (Piper et al., 2011; Simpson et al., 2017). These data indicate that lifespan and reproduction are not in competition for limiting energy derived from the diet, but instead are optimised at different dietary protein : carbohydrate ratios. In response to these findings, an enormous effort is now focused on uncovering how macronutrient rebalancing, in particular protein dilution, acts to improve lifespan (Blagosklonny, 2006, 2010; Moatt et al., 2020; Regan et al., 2020; Speakman, 2020). Accumulating evidence indicates that the effect is mediated by reducing signalling through the amino acid sensitive Target Of Rapamycin (TOR) pathway to enhance cellular proteostasis (Sanz *et al.*, 2004; Ayala *et al.*, 2007; Raubenheimer & Simpson, 2009; Simpson & Raubenheimer, 2009; Taylor & Dillin, 2011; Fansonet *et al*, 2012; Sabatini, 2017).

Although detrimental for lifespan, relatively high protein, low carbohydrate diets are beneficial for female reproduction (Chong *et al.*, 2004; Solon-Biet., *et al.*, 2015). We have studied this closely in the fruitfly *Drosophila melanogaster*, where the principle driver of egg production is dietary protein (Min & Tatar, 2006; Grandison *et al.*, 2009; Piper et al., 2017). Although protein is key, females must transfer dozens of nutrients into eggs for future embryo formation and not all of these components contribute to the flies’ decision to produce eggs (Piper *et al.*, 2014; Mirth *et al.*, 2019; Wu *et al.*, 2020). This means that high protein diets might drive mothers to produce eggs at a faster rate than they can support if the diet contains insufficient levels of the other components that are required to make eggs. In this scenario, the macronutrients would have an indirect effect on lifespan by changing the availability of another limiting nutrient for somatic maintenance. If true, this would move the focus of mechanistic studies away from the direct effects of protein, TOR and proteostasis, towards some other component of nutritional physiology. Distinguishing between these possible causes of death is important since it would fundamentally change our understanding of the way diet alters lifespan. It also has the important knock-on effect that we could change the way we design diets for longer life. For instance, supplementing high protein diets with key limiting nutrients would be as beneficial as restricting dietary protein or treating with pharmacological suppressors of TOR.

Of the many studies that have examined the effects of dietary protein and carbohydrate on lifespan and reproduction in *Drosophila*, most have done so by varying dietary yeast and sugar proportions, where yeast is the flies’ natural source of protein (Mair *et al.*, 2005; Lee *et al.*, 2008; Skorupa *et al.*, 2008). However, yeast also contains all of the fly’s other essential macro and micronutrients whose relative proportions can change, and thus possibly interact with protein and carbohydrates to dictate life history outcomes. We have previously found that depriving adult female flies of a source of sterols, an essential micronutrient for insects, imposes a minor cost on reproduction, but a substantial (>50%) cost to lifespan (Piper *et al.*, 2014; Wu *et al.*, 2020). These data indicate that yeast sterol levels may contribute to the effects on lifespan of protein and carbohydrate. To investigate the interactions between dietary protein, carbohydrate, and sterols systematically, we have used the design principles of the geometric framework for nutrition (Simpson & Raubenheimer, 2012; Simpson & Raubenheimer, 1993) and a completely defined (holidic) diet that allows us to control the levels of each nutrient independently of all others (Piper *et al.*, 2014; Piper *et al.*, 2017). These data point to an important role for sterols in determining *Drosophila* lifespan, which we verified to be relevant in two yeast based media that are often used in *Drosophila* lifespan studies. This work is critical to identifying how diet modifies lifespan at the molecular level, and highlights a new way to think about diet design to improve healthy ageing.

## Results

### Protein: carbohydrate ratio influences lifespan and reproduction

To examine the interactive effects of dietary protein, carbohydrate, and cholesterol on *Drosophila* lifespan and fecundity, we used our completely defined (holidic) diet (Piper *et al.*, 2014) to manipulate each nutrient independently of all others. We selected dietary protein and carbohydrate concentrations that we know to elicit the full range of lifespan and fecundity responses to dietary restriction (Lee *et al.*, 2008; Piper *et al.*, 2014, 2017; Ma *et al.*, 2020).

Similar to what we and others have found previously (Mair *et al.*, 2005; Lee *et al.*, 2008; Grandison *et al.*, 2009; Piper *et al.*, 2014, 2017; Katewa *et al.*, 2016), lifespan and reproduction were modified by dietary protein manipulations (Figure 1). Specifically, egg production showed a linear, positive correlation with dietary protein content (Supplementary table 3), while lifespan showed a peak at intermediate protein (67 d median at 10.7 g/l), and fell away at both higher (48d median at 33.1 g/l) and lower (43 d median at 5.2 g/l) concentrations (Figure 1 a-b; Supplementary Table 4). Thus, as is typical for dietary restriction experiments, restricting dietary protein from high to intermediate levels increased lifespan and decreased reproduction (Lee *et al.*, 2008; Skorupa *et al.*, 2008; Grandison *et al.*, 2009; Katewa *et al.*, 2016; Le Couteur *et al.*, 2016).

**Figure 1.**
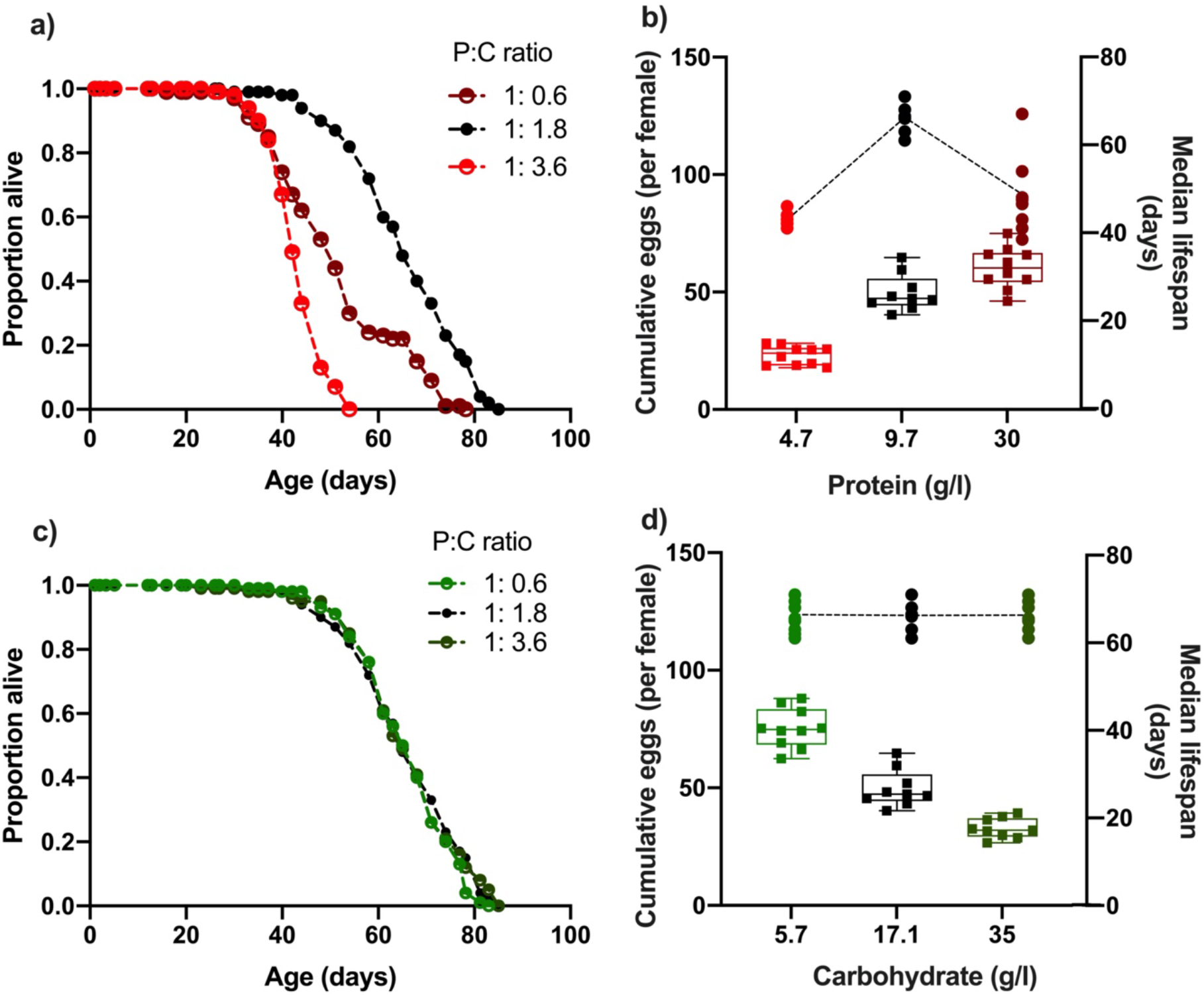
Changing dietary protein and carbohydrate concentrations modify *Drosophila* lifespan and fecundity. (a, b) Lifespan was maximised at our intermediate dose of dietary protein (carbohydrate fixed at 17.1 g/l) but was unaffected by our carbohydrate (c, d) concentration range (protein fixed at 10.7 g/l). (b) Cumulative egg production had a significant positive relationship with protein levels and (d) significant negative correlation with dietary carbohydrate content. Note that the intermediate protein and carbohydrate diet (10.7g/l protein, 17.1g/l carbohydrate) is common to both nutrient dilution series. The median survival data in panels (b) and (d) represent data from replicates that are combined in panels (a) and (c) respectively. Statistical analyses reported in Supplementary Table 3 and Supplementary Table 4.

When increasing dietary carbohydrate against an otherwise fixed nutritional background, egg laying was suppressed in a dose-dependent fashion, but lifespan remained at its maximum level and was unchanged across all carbohydrate doses (~66d median, Figure 1 c-d). The diet with the lowest concentration of carbohydrate (5.7g/l), which also contained the intermediate protein level (10.7g/l), supported both maximum lifespan (Figure 1d; 67 d median) and the highest level of egg laying (75 eggs/female) of any diet in our experiment. Thus, as we have previously shown (Piper *et al.*, 2017), balancing the dietary protein and carbohydrate concentrations can reveal a single dietary optimum for both traits, showing that lifespan shortening is not necessarily caused by high egg laying alone.

### Cholesterol interacts with protein and carbohydrate to modify lifespan and reproduction

Most dietary restriction studies on *Drosophila* vary dietary protein by modifying the yeast levels in food (Chapman & Partridge, 1996; Mair *et al.*, 2005; Lee *et al.*, 2008; Skorupa *et al.*, 2008). While yeast is the flies’ major source of protein, it is also their only source of dozens of other nutrients, including sterols, which are essential micronutrients for insects (Carvalho et al., 2010). To quantify the effects of varying dietary sterol levels on fly lifespan and egg laying, we maintained flies on the same set of diets as above, varying in protein and carbohydrate concentrations, while also varying cholesterol across four different levels: 0 g/l, 0.15g/l (low), 0.3 g/l (medium; also our standard level) and 0.6 g/l (high).

We first compared the flies’ responses to variation in both protein and cholesterol (Figure 2). In general, lifespan was optimised at our intermediate dose of protein, while increasing cholesterol was beneficial, but with diminishing effects as its concentration was increased (Supplementary Figure 1 a-b and Supplementary Table 5). Interestingly, changing cholesterol modified the flies’ lifespan response to protein, an effect that can be seen when the data are separated by level of cholesterol addition (Figure 2). At 0 g/l cholesterol (Figure 2a) increasing protein concentration in the diet decreased lifespan. However, at 0.15 g/l cholesterol, the shape of the response changed such that only the highest protein concentration decreased lifespan (35d median; Figure 2b) when compared with intermediate (9.7 g/l; 55d median) and low protein (4.7 g/l; 52d median) diets. At 0.3 g/l of cholesterol, lifespan was highest on the diet with intermediate protein concentration (66d median) and flies on the high protein diet were longer lived (49d median) than the flies on the lowest protein diet (43d median). Finally, increasing cholesterol from 0.3 g/l to 0.6 g/l (Figure 2d) did not change the way that lifespan responded to protein. Thus, lowering dietary cholesterol was detrimental for lifespan and it intensified the negative effects of increasing dietary protein concentrations.

**Figure 2.**
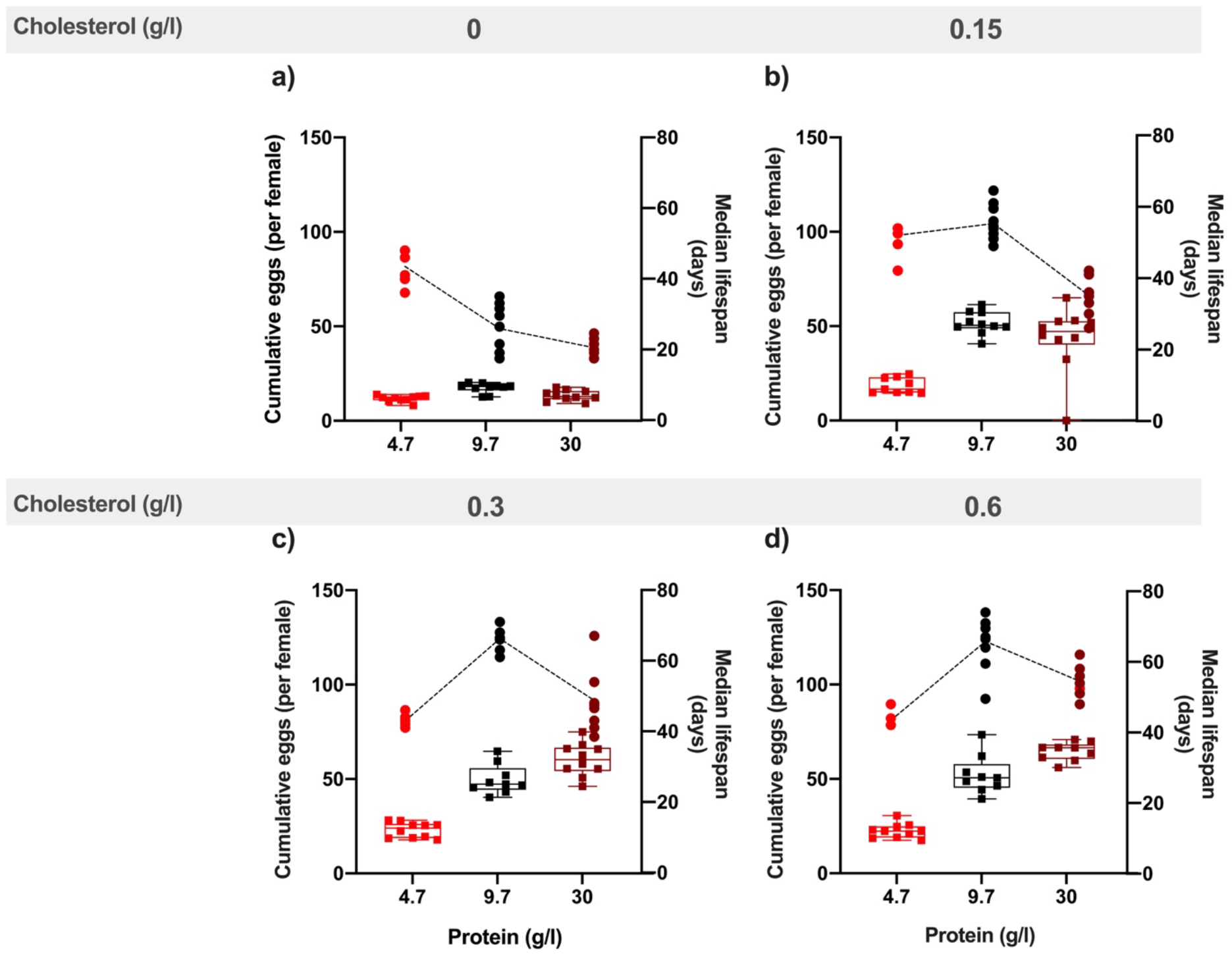
Dietary cholesterol content significantly modified the effect of protein content on lifespan and reproduction. Lowering cholesterol most severely compromised lifespan as protein levels increased. In general, increasing protein drove increasing levels of egg production and this was enhanced by increasing dietary cholesterol levels. Cumulative eggs laid per female are shown as box plots. Lines show the relationship between dietary protein and median survival (days) for each cholesterol level. (0g/l (a), 0.15g/l (b), 0.3g/l (c), 0.6g/l (d)). Statistical analyses reported in Supplementary Table 5 and Supplementary Table 6.

Across the same set of diets, we observed a generally beneficial effect on egg laying of increasing dietary protein and cholesterol, and both had diminishing benefits as their concentrations increased (Supplementary Figure 1 d-e; Supplementary Table 6). Cholesterol also modified the way egg laying was affected by dietary protein (Figure 2). Increasing cholesterol from 0g/l (Figure 2a) to 0.15 g/l (Figure 2b) amplified the positive effect on egg laying of increasing dietary protein. Further increasing cholesterol to 0.3 g/l had an additional benefit for egg laying (Figure 2c), but only for flies on the highest protein diet (compare Figure 2b with Figure 2c), while increasing cholesterol even further, to 0.6 g/l (Figure 2d), did not change egg laying from that seen on 0.3 g/l. Thus, the response of egg laying to increasing protein was only compromised when cholesterol was completely removed from the diet, or when cholesterol was low (0.15 g/l) and protein was high (30 g/l) (Figure 2b).

Together, these data show that reducing cholesterol had negative effects on both lifespan and egg laying, and that these negative effects became more pronounced with increasing dietary protein. Furthermore, the negative interaction between lowering cholesterol and increasing protein was more severe and occurred at a lower protein concentration for lifespan than it did for egg laying.

Next, we looked to see if changing dietary cholesterol modified the responses of lifespan and egg laying to variation in carbohydrate concentration (Supplementary Figure 1 c & f; Supplementary Table 5). At 0 g/l cholesterol, lifespan was generally short (31d median) but positively affected by increasing dietary carbohydrate (up to 40 d median) (Figure 3a). As dietary cholesterol was increased to 0.15 g/l, lifespan on all diets was higher and the positive effect of increasing carbohydrate was preserved (Figure 3b). However, when cholesterol reached 0.3 g/l, the flies were constantly long-lived, and lifespan was unaffected by dietary carbohydrate level (66d median) (Figure 3c). This pattern was not changed by increasing cholesterol further to 0.6 g/l (Figure 3d). Thus, each of our dietary carbohydrate levels could support maximal fly lifespan, but the lower carbohydrate diets were more susceptible to the detrimental effects of dietary cholesterol dilution.

**Figure 3.**
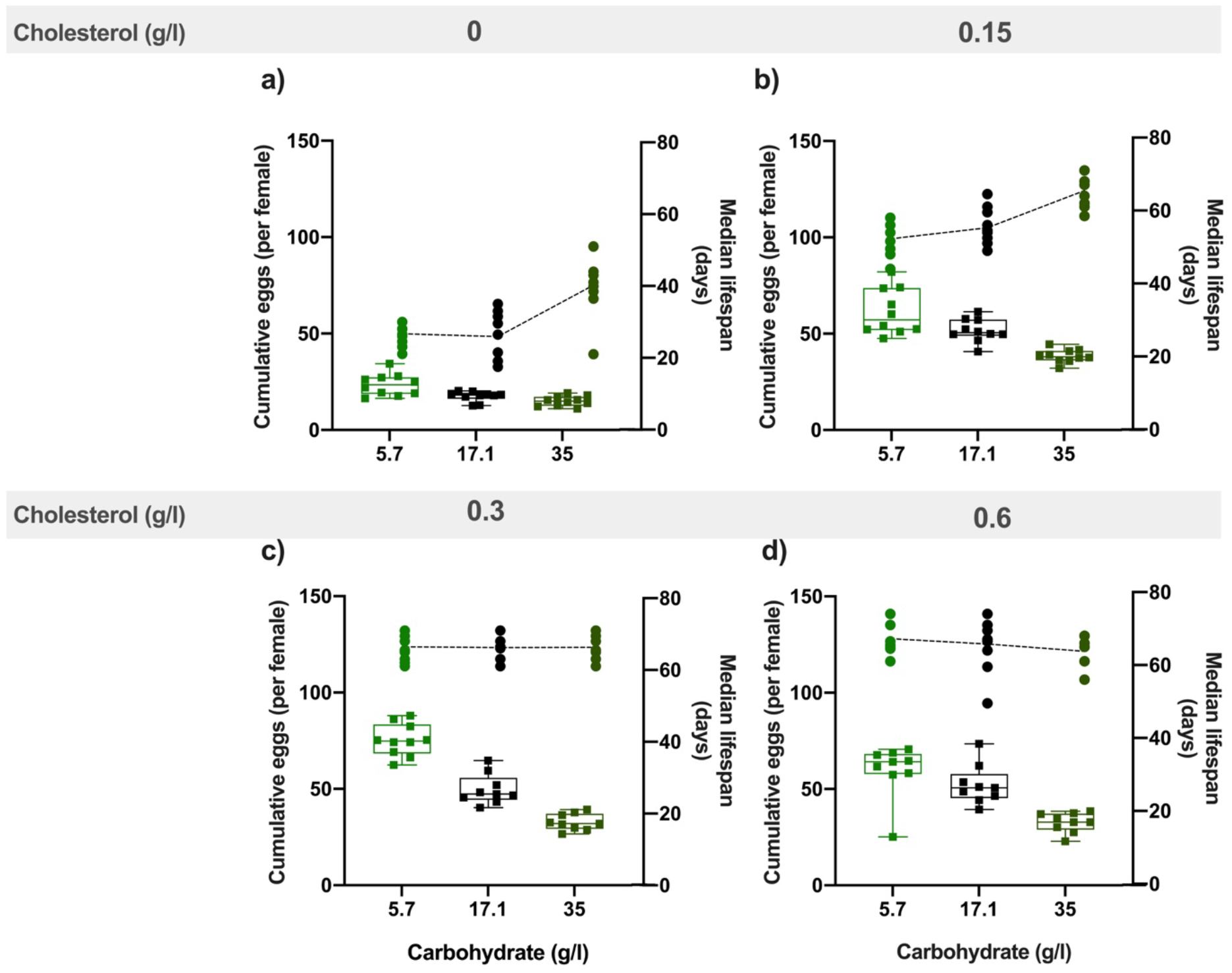
Dietary cholesterol content significantly modified the effect of carbohydrate content on lifespan and on reproduction. Lowering cholesterol most severely compromised lifespan as carbohydrate levels decreased – the conditions that drove increasing levels of egg production. Cumulative eggs laid per female are shown as box plots. Lines show the relationship between dietary carbohydrate and median survival (days) for each cholesterol level. (0g/l (a), 0.15g/l (b), 0.3g/l (c), 0.6g/l (d)). Statistical analyses reported in Supplementary Table 5 and Supplementary Table 6.

Increasing dietary carbohydrate had a generally negative impact on egg laying and this effect was modified by the benefits of increasing dietary cholesterol (Figure 3; Supplementary Figure 1f; Supplementary Table 6). Without any cholesterol in the food, egg laying was consistently low and was negatively affected by increasing dietary carbohydrate (Figure 3a). This negative effect of carbohydrate on egg laying became stronger as cholesterol was increased to 0.15 g/l (Figure 3b) and 0.3 g/l (Figure 3c), with no further change as cholesterol increased from 0.3 g/l to 0.6 g/l (Figure 3d). This increasingly negative relationship between carbohydrate and egg laying was caused because increasing cholesterol benefited egg laying more at lower dietary carbohydrate levels – the opposite of what we observed for lifespan.

Thus, once again fly lifespan and egg laying worsened as cholesterol was diluted, but unlike its negative interaction with *increasing* dietary protein, the detrimental effects of lowering cholesterol became stronger as carbohydrate levels *decreased.* This indicates that the negative impact of lowering cholesterol is not a specific interaction with either high protein or low carbohydrate levels in the diet. Instead lowering cholesterol produces worse outcomes as the dietary protein : carbohydrate ratio increases. This is the same change in macronutrient balance that promotes increasing egg laying.

### Increasing the dietary protein : carbohydrate ratio drives increasing reproduction and makes fly lifespan susceptible to dietary cholesterol dilution

We saw that flies produce more eggs in response to increasing dietary protein : carbohydrate ratio and that these positive effects persist even as dietary cholesterol falls to a level that cannot fully support lifespan (less than 0.3 g/l cholesterol). Thus, the protein : carbohydrate ratio appears to take precedence over dietary sterol levels in the decision to commit to reproduction. If this is the case, we expect to see a positive relationship between the dietary protein : carbohydrate ratio and egg laying across all levels of dietary cholesterol. This is indeed what we found, although the positive relationship was modified by cholesterol level, starting with a weak positive effect on 0 g/l cholesterol (Figure 4a) and becoming increasingly positive as cholesterol was increased to 0.15 g/l (Figure 4b) and 0.3 g/l (Figure 4c). Once again, increasing cholesterol from 0.3 g/l to 0.6 g/l promoted no further benefit (Figure 4d, Supplementary Table 7).

**Figure 4.**
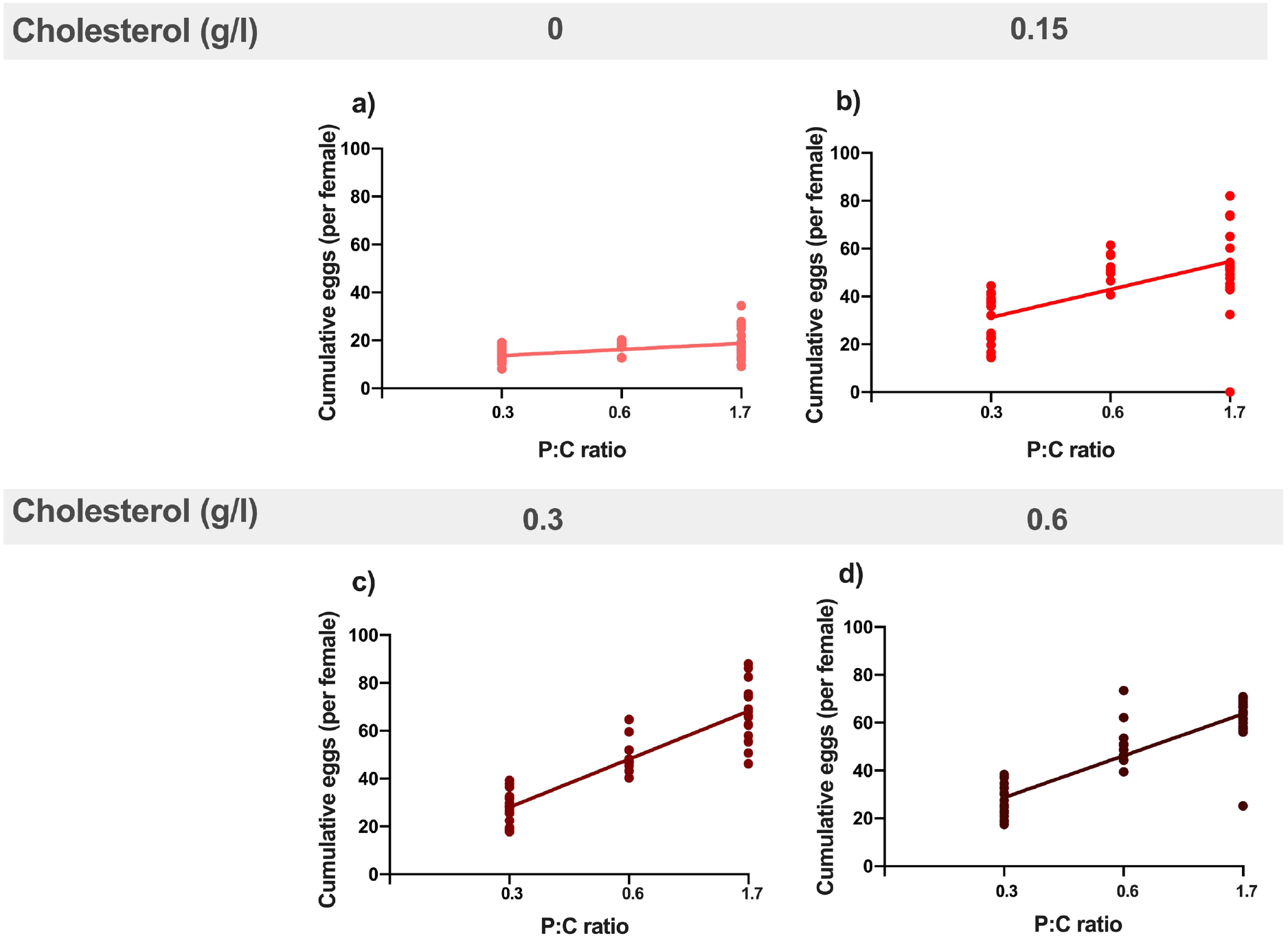
Increasing dietary protein : carbohydrate ratio resulted in increased egg production at every level of cholesterol. This positive effect was stronger from 0g/l cholesterol (a) to 0.15 g/l (b) and 0.3 g/l (c). There was no additional benefit of further increasing cholesterol to 0.6 g/l (d). Regression lines show the relationship between cumulative eggs laid per female and protein : carbohydrate ratio. Statistical analysis reported in Supplementary Table 7.

Reproduction can impose a cost on future survival if resources that are essential for somatic maintenance are preferentially committed to making eggs. Since increasing protein : carbohydrate levels drove increasing egg laying, even when the adults were completely deprived of sterols, it is possible that females are committing sterols to egg production at a rate faster than they can replenish it from the diet. If true, mothers on low cholesterol diets would become shorter lived as egg laying increases, but when cholesterol is sufficient, the relationship between egg production and lifespan should become less negative. To test this, we plotted egg laying against lifespan for all replicates across all diets. This showed that egg laying was a significant predictor of lifespan, and that this relationship was modified by dietary cholesterol (Supplementary Table 8). When the data are grouped by dietary cholesterol level (Figure 5), we see that when cholesterol was at 0g/l (Figure 5a), there was a negative relationship between egg laying and lifespan, but as the cholesterol level increased, the correlation flattened, such that the slope was no longer negative for each level of cholesterol supplementation (Figure 5b-d, Supplementary Table 8).

**Figure 5.**
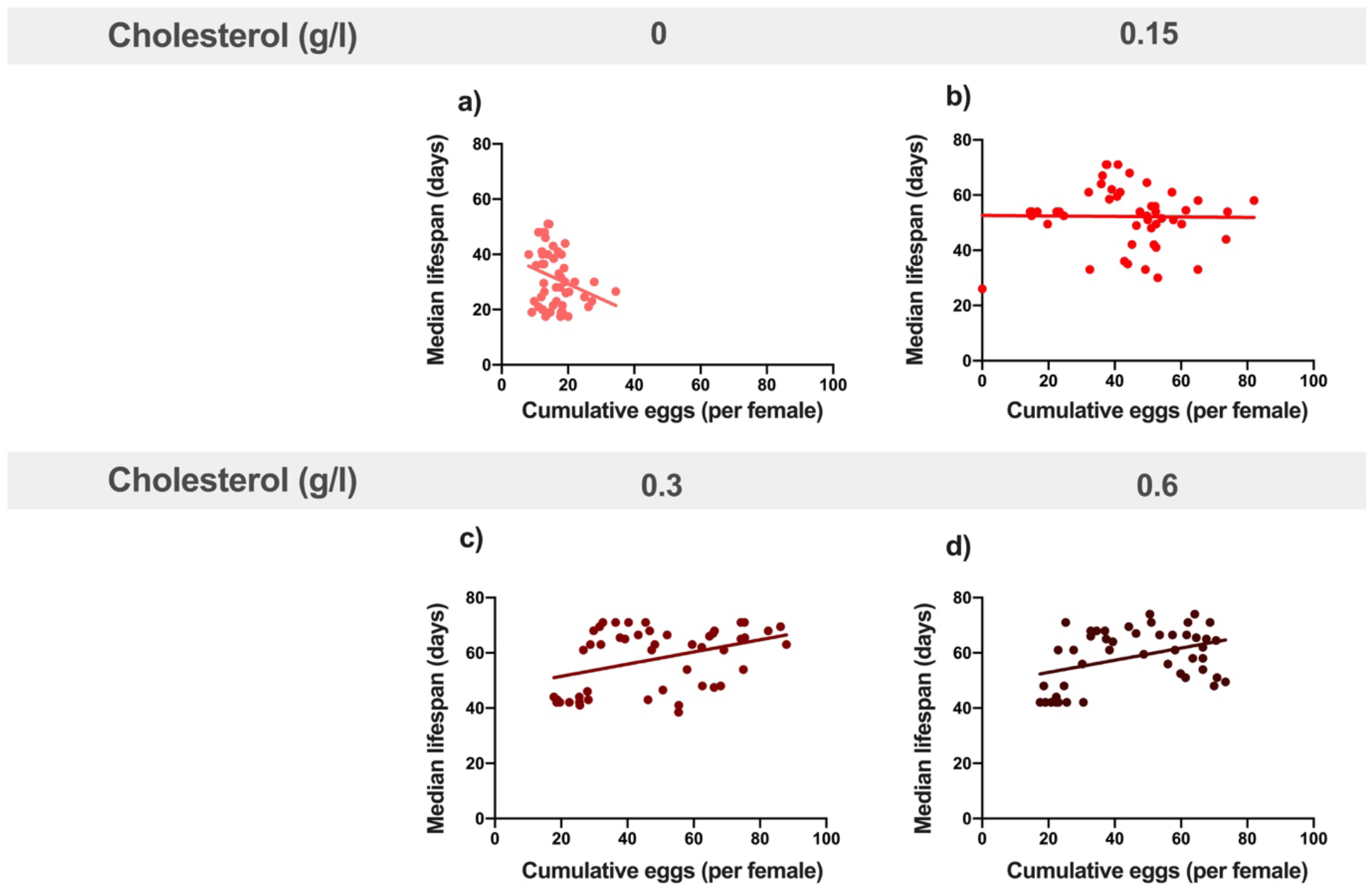
Providing adequate cholesterol transforms the relationship between egg production and lifespan from negative to positive. When cholesterol was not available (0g/l) (a), there was a negative relationship between egg laying and lifespan as dietary protein and carbohydrate levels were varied. When cholesterol was provided at (0.15g/l)(b) or above (c, d), this negative relationship was eliminated and egg production varied independently of lifespan. Regression lines show the relationship between cumulative eggs laid per female and median survival (days) in response to varying levels of cholesterol availability. Statistical analysis reported in Supplementary Table 8.

Thus, when dietary cholesterol was insufficient, increasing dietary protein : carbohydrate drove higher egg laying (Figure 4a) and this predicted lifespan shortening (Figure 5a) – a scenario that exemplifies the negative relationship between reproduction and lifespan in response to increasing protein : carbohydrate levels that is regularly reported in the dietary restriction literature (Mair *et al*., 2005; Lee *et al.*, 2008; Skorupa *et al.*, 2008; Solon-Biet *et al.*, 2014; Solon-Biet *et al.*, 2015; Simpson *et al.*, 2017). However, when cholesterol was increased, the negative relationship was reduced such that egg laying was either completely independent of lifespan (Figure 5b) or became slightly positively correlated, indicating that the dietary conditions which promote egg laying are the same as those that promote longer lifespan (Figure 5c-d). Thus, higher egg laying in response to increasing protein : carbohydrate levels only shortens lifespan when cholesterol is insufficient to support egg production.

### Lifespan extension by rapamycin depends on dietary cholesterol level

TOR signalling is a key amino acid signalling pathway that is critical for growth, reproduction and lifespan. Because TOR activity increases with dietary protein levels, it has been implicated as mediating the detrimental effects on lifespan of high protein diets (Liu & Sabatini, 2020). This is supported by the fact that rapamycin, a pharmacological suppressor of TOR, has been shown to extend lifespan across numerous species, including *Drosophila* where it also suppresses egg production (Bjedov et al., 2010; Schinaman, Rana, Ja, Clark, & Walker, 2019; Scialò et al., 2015). If cholesterol limitation is the reason why high egg production on high protein : carbohydrate diets causes reduced lifespan, rapamycin might extend lifespan because it reduces egg production and therefore rescues females from cholesterol depletion. If true, rapamycin should extend life only when the flies on high protein : carbohydrate diets are cholesterol limited.

As before, when we maintained flies on a high protein : carbohydrate diet, increasing dietary cholesterol from 0.1 g/l to 0.3 g/l increased lifespan (62 d median v 69 d median)(Figure 6a). Egg laying was also slightly (34%), but significantly, elevated by cholesterol supplementation (Figure 6b) indicating that 0.1 g/l cholesterol was limiting for both lifespan and reproduction. When rapamycin was added to both foods, egg laying was almost completely suppressed (Figure 6b). Rapamycin also extended fly lifespan, but only for flies on low dietary cholesterol (0.1 g/l)(Figure 6a), bringing their lifespan up to the same level as flies on higher cholesterol food (0.3 g/l; 69 d median). Adding rapamycin to the food with higher cholesterol did not result in any additional lifespan improvement over what was already achieved by increasing cholesterol alone (69 d median; Figure 6a).

**Figure 6.**
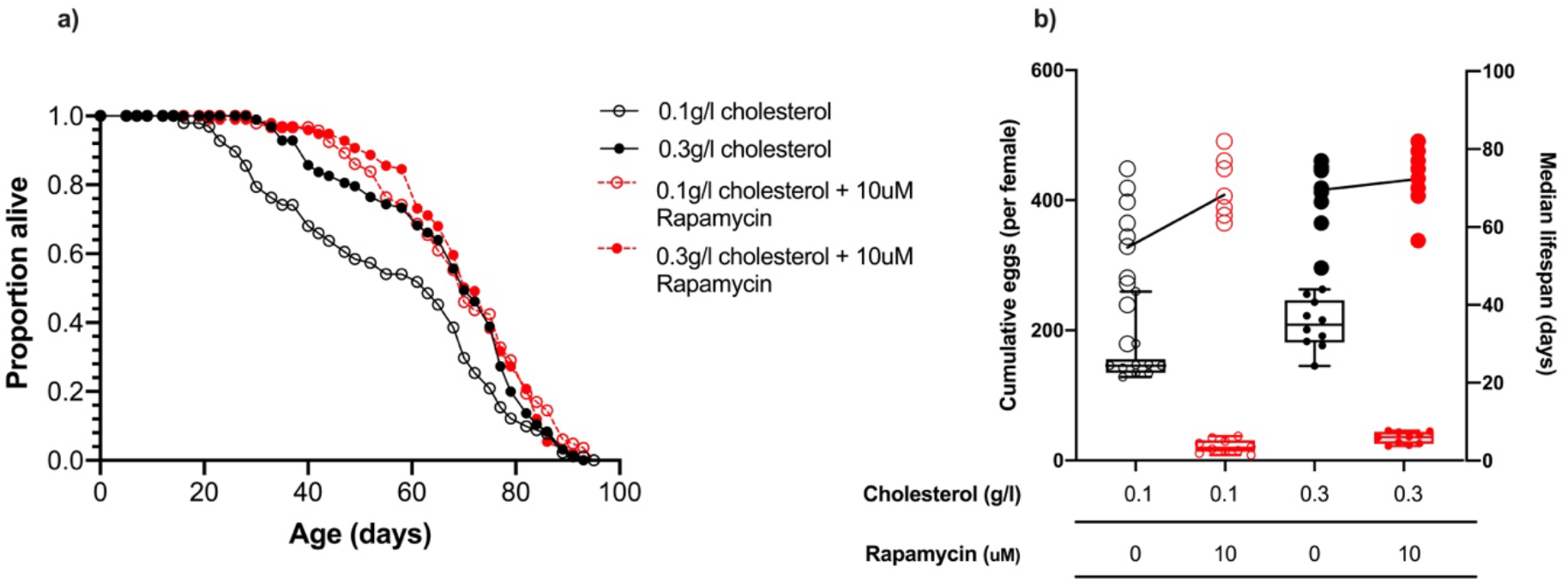
Rapamycin extends lifespan in flies consuming a low cholesterol diet (0.1 g/l) but had no effect when cholesterol level was increased to 0.3 g/l. (a) There was no significant difference in lifespan among flies fed 0.3g/l cholesterol, 0.3g/l cholesterol + rapamycin or 0.1mg/l cholesterol + rapamycin, all of which were significantly longer lived than flies fed 0.1g/l cholesterol (0.1g/ v 0.3g/l, *P* = 0.014; 0.1g/l v 0.1g/l + rapamycin*, P* < 0.001; 0.1g/l v 0.3g/l + rapamycin, *P* = 0.002, log rank test). (b) Cumulative eggs laid were significantly reduced in flies treated with rapamycin (*P* < 0.001, ANOVA), and also significantly reduced when cholesterol was limited (p<0.001, ANOVA).

These data show that lifespan extension by rapamycin administration is conditional on the flies being on a low cholesterol diet. Together, our data are consistent with the fly’s lifespan being determined by having access to sufficiently high levels of dietary sterols that they have enough left over after reproduction to meet their needs for somatic maintenance. This can be achieved either by enriching the amount of cholesterol in the diet, or by reducing the flies’ expenditure on egg production, which can be achieved by reducing the dietary protein : carbohydrate ratio or by suppressing egg production pharmacologically.

### Standard yeast-based media used in the laboratory contains lifespan limiting levels of sterols

The experiments above were all performed using synthetic diets in which our ability to vary the absolute and relative concentrations of protein, carbohydrate, and sterol are limited only by solubility. However, most laboratories maintain fly populations on a diet that consists of yeast and sugar as well as variable numbers of other ingredients (Piper, 2017). Although the relative concentration of each nutrient in yeast is more constrained than on our synthetic diet, systematic studies have shown that the type and commercial source of yeast can have significant effects on overall dietary composition (Lesperance & Broderick, 2020) and the relationship between lifespan and egg laying (Bass et al, 2007). In Bass *et al.* (2007), the most dramatic lifespan reduction for increasing yeast was found when the fly food was made with a water-soluble extract of yeast that would contain very little, if any, sterols. Thus, similar to what we demonstrated on the synthetic diet, the shortening of fly lifespan when increasing the yeast content (protein : carbohydrate ratio) in lab foods may be caused by an insufficiency of dietary sterols.

We tested the effects of supplementing cholesterol into two sugar / yeast recipes that have been commonly used to study the effects of dietary restriction on lifespan (Mair *et al.*, 2005; Bass *et al.*, 2007; Katewa *et al.*, 2016). These diets differ in both the number of ingredients used and the type of yeast; while both are *Saccharomyces cerevisiae*, one is a whole cell autolysate, while the other is a water-soluble extract. Adding 0.3 g/l cholesterol to both the low yeast (dietary restriction) and high yeast foods of both yeast types had a significant positive effect on lifespan (Figure 7a, c) and egg laying (Figure 7b, d) when compared to diets without cholesterol supplementation. The magnitude of this benefit to lifespan was greater for flies on the high yeast foods than on the low yeast foods, meaning that cholesterol supplementation narrowed the difference between the dietary restriction vs high yeast diet from 9% to 4% for flies on the autolysed yeast diets (Figure 7a) and from 81% to 25% lifespan extension for flies on the yeast extract diets (Figure 7c). We note that even with cholesterol supplementation, the flies on the high yeast diet were still significantly shorter lived than those on the cholesterol supplemented low yeast food. This small additional cost of the high yeast food could reflect a detrimental (toxic) effect on lifespan of very high dietary protein, similar to what we observed in our highest protein diets on the synthetic foods (Figure 2 c, d). This is not rescuable by cholesterol supplementation and is not related to the number of eggs that females produce.

**Figure 7.**
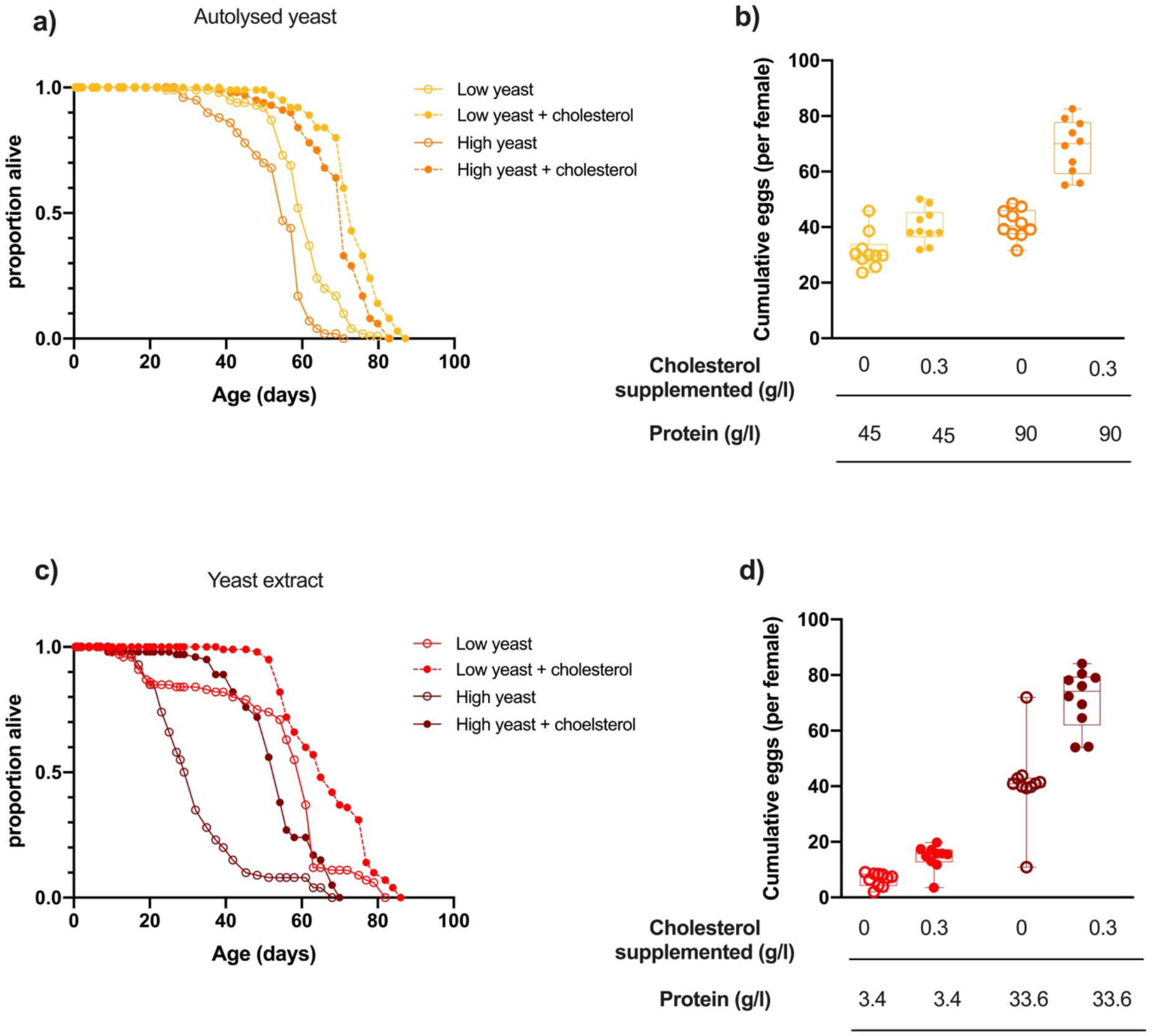
Cholesterol supplementation significantly extended lifespan and promoted egg laying of flies fed yeast-based diets. (a) Adding dietary cholesterol significantly increased the lifespan of flies on both high and low concentrations of diets made with autolysed yeast (low yeast v low yeast+cholesterol and high yeast v high yeast + cholesterol; *P* < 0.001, log rank test). (b) Yeast and cholesterol addition to these two foods both positively affected egg production (*P* <0.001, two-way ANOVA). (c) Cholesterol addition significantly extended the lifespan of flies on diets made with yeast extract (low YE v low YE+cholesterol and high YE v high YE + cholesterol; *P* <0.001, log rank test). (d) Cumulative egg laying was also positively affected by yeast addition and cholesterol addition to each yeast level (P <0.001, two-way ANOVA).

## Discussion

The reason that higher protein : carbohydrate diets shorten lifespan in DR studies is routinely attributed to their direct effects on nutrient signalling pathways and physiology. However our data implicate a fundamentally different mechanism, in which the macronutrients act indirectly, by manipulating sterol availability, which then modifies lifespan. Specifically, diets with high protein : carbohydrate ratios decrease lifespan by causing mothers to overinvest limiting sterols into egg production. Thus, although the macronutrients set egg laying rates, it is actually the sterols that determine lifespan due to a trade-off with reproduction. The corollary of this finding is that the lifespan of flies on high protein : carbohydrate diets can be extended by increasing the supply of cholesterol. This approach is the opposite of, but complementary to, the already recognised strategies to extend lifespan by DR, which reduce maternal investment into reproduction by decreasing the dietary protein : carbohydrate ratio (Mair *et al.*, 2005; Lee *et al.*, 2008; Skorupa *et al.*, 2008) or by treating the animals with rapamycin that suppresses TOR and reduces reproduction (Bjedov et al., 2010; Harrison et al., 2009; Liu & Sabatini, 2020). It is also consistent with our recent work that showed non-reproducing adult males and genetically sterile females suffer little to no lifespan cost when sterol deprived, which is presumably because they conserve sterols which would otherwise be depleted by reproduction (Wu et al, 2020).

### High protein diets promote egg production, driving a lethal micronutrient deficiency

In the lab, flies can be successfully reared and maintained on a mixture of just sugar and yeast (Pearl & Parker, 1921). This diet is thought to reflect their natural diet of rotting fruit and the microbial community – principally the yeasts – that cause the fruit to decay (Markow *et al.*, 2015; Piper, 2017). Yeast contains all of the nutrients that flies require, including protein (~45%), carbohydrate (~40%), a small amount of fat (~7%), nucleic acids (~7%), and micronutrients, such as sterols, metal ions and vitamins, which are essential for flies. *Drosophila* rely heavily on protein from yeast, as well as carbohydrate from both yeast and plant sources, to guide their feeding behaviour. They select amongst foods containing the appropriate protein and carbohydrate concentrations to enhance their fitness (Ribeiro & Dickson, 2010; Vargas *et al.*, 2010; Walker *et al.*, 2017). Many of the other nutrients from their diet, including sterols, do not affect feeding behaviour, presumably because they are normally acquired in adequate quantities as part of a diet that is sufficient in macronutrients (Walker, *et al.*, 2015; Leitão-Gonçalves *et al.*, 2017; Münch *et al.*, 2020).

While the relative proportion of protein and carbohydrate in yeast remains relatively constant across growth conditions, the abundance of sterols can vary over a 10-fold range in response to changes in oxygen availability, which is essential for sterol biosynthesis (Starr & Parks, 1962; Wilson & McLeod, 1976). Thus, because fly feeding behaviour and egg production are almost entirely shaped by the macronutrients, fly lifespan is susceptible to reductions in the sterol : protein content of dietary yeast. Our data indicate that this is because protein drives sterols to be preferentially partitioned towards reproduction at the expense of maintaining the adult soma. While we have found this to be the case for flies feeding on lab based foods, it is also reasonable to expect it for flies feeding on rotting fruit, where microbial growth is largely fermentative (driven by high sugar levels and limiting oxygen), producing ethanol and short chain acids to which *Drosophila* has evolved a healthy tolerance (Geer *et al.*, 1993).

### Extending fly lifespan by DR involves an indirect trade-off

There have been several theoretical attempts to describe the mechanistic basis for the lifespan benefits of dietary restriction (Blagosklonny, 2006, 2010; Kirkwood & Rose, 1991; Moatt et al., 2020; Regan et al., 2020; Speakman, 2020). In particular, the disposable soma theory proposes that organisms will strategically reallocate nutrients towards somatic maintenance at the cost of reproduction when nutrients are scarce and that this enhances lifespan (Kirkwood & Rose, 1991). Our data indicate that this trade-off can exist for flies feeding on yeast, but only when dietary sterols are limiting. However, when dietary sterols are not limiting, this trade-off does not need to exist and a single nutritional optimum for both lifespan and reproduction can be found. Thus, the macronutrient balance that drives higher egg laying does not necessarily inflict a direct cost on lifespan.

In mechanistic work, the increased lifespan under dietary restriction has been attributed to the benefits of reduced dietary protein, which enhances proteome maintenance via reduced TOR signalling (Harrison *et al.*, 2009; Partridge *et al.*, 2011; Kapahi *et al.*, 2017; Piper *et al.*, 2017; Sabatini, 2017; Dobson *et al.*, 2018; Liu & Sabatini, 2020). However, we have shown that flies on a high protein : carbohydrate diet, in which TOR signalling would be expected to remain high, can still sustain a long lifespan if supplemented with cholesterol, or alternatively, if the cost of reproduction on sterol stores is removed by making flies infertile (Wu et al, 2020). This demonstrates that the major improvement to lifespan observed on low protein diets is not the result of enhanced proteostasis, but is instead, a side effect of avoiding sterol depletion caused by egg production. It is likely that a deficiency of this essential micronutrient could compromise the integrity of critical tissues, such as the gut, whose quality is strongly associated with lifespan changes under dietary restriction (Rera *et al.*, 2013; Regan *et al.*, 2016).

We note that while dietary sterols can account for the dietary restriction effect seen for flies feeding on yeast, other limiting micronutrients could be key for explaining the lifespan benefits of dietary restriction in other organisms. If true, supplementing their high food diets with limiting micronutrient(s) should mimic the benefits of dietary macronutrient remodelling as sterol supplementation did for flies in our experiments.

### Conclusion

Our data show that the detrimental effects of a high protein : carbohydrate diet on lifespan in female *Drosophila melanogaster* are, to a significant extent, driven by an indirect nutrient trade-off, in which the macronutrients drive maternal sterol depletion by enhancing egg laying. This is a fundamentally different mechanism from the predominant view that reducing protein : carbohydrate balance in diets improves lifespan by a direct action to reduce TOR signalling and enhance proteostasis. Because of our discovery, we show that the shortened lifespan of flies on a high protein : carbohydrate diet can be improved by supplementing their diet with cholesterol, as well as by reducing egg production by lowering the dietary protein : carbohydrate ratio or by administering rapamycin. Further work is now needed to discover the mechanisms through which cholesterol works to modify lifespan in *Drosophila melanogaster*, and the role of other important micronutrients in healthy ageing across taxa.

## Methodology

### Fly husbandry

All experiments were conducted using a wild type *Drosophila melanogaster* strain called Dahomey (Mair *et al.*, 2005). These flies have been maintained in large numbers with overlapping generations to maintain genetic diversity. Upon removal from their population cages, flies were reared for two generations at a controlled density before use in experiments, to control for possible parental effects. Eggs for age-synchronised flies were collected over 18 hours, and the resulting adult flies emerged during a 12-hr window. They were then allowed to mate for 48 hr before being anaesthetised with CO2, at which point females were separated and allocated into experimental vials. Stocks were maintained and experiments were conducted at 25°C on a 12 hr:12 hr light:dark cycle at 65% humidity (Bass et al., 2007).

### Lifespan assays

For all lifespan assays, flies were placed into vials (FS32, Pathtech) containing 3ml of treatment food at a density of ten flies per vial, with ten replicate vials per treatment. Flies were transferred to fresh vials every two to three days at which point deaths and censors were recorded and saved using the software package Dlife (Linford *et al*., 2013; Piper & Partridge, 2016).

### Fecundity assays

Fecundity was measured as the sum of the mean number of eggs laid per female once per week over four weeks (commencing on day 8 of the experiment), except for the sugar yeast (SY) medium experiment, for which egg counts were recorded in weeks one, two and three. These timepoints were selected because measuring reproductive output during the first weeks of egg laying has shown to be representative of life-long fecundity in flies (Chapman & Partridge, 1996). The eggs laid on the food surfaces of all vials were imaged using a web camera mounted on a Zeiss dissecting microscope and eggs were counted both manually and using Quantifly (Waithe *et al.*, 2015). Quantifly was trained using five images for each cholesterol concentration due to variance in food opacity.

### Experimental Diets

#### Holidic medium experiments

To examine the effects of protein: carbohydrate ratio on lifespan and fecundity we chose five experimental diets that consisted of three different protein (amino acid): carbohydrate (sucrose) ratios at three levels of similar caloric densities (Figure 1a, Table 1). These diets also made up a three-diet series of protein only dilution, and a three-diet series of carbohydrate only dilution (Figure 1a). The two diet series had one diet in common, which was our most commonly used, “standard” lab diet (Piper et al, 2014). These diets incorporate those known to maximise either lifespan, reproduction or both (Ma et al., 2020; Piper et al., 2017). To examine the effects of cholesterol on these traits, we selected four cholesterol concentrations for each of these five diets, making a total of 20 diets (Figure 1b, Table 1). All diets were made using the holidic medium described in Piper et al. (2014), in which free amino acids are used to make up protein equivalents. To convert amino acids to protein equivalents, we used the molar quantities of nitrogen and the assumption that N makes up 16% of whole proteins (Sosulski and Imafidon, 1990). In this case, an amino acid ratio matched to the exome of adult flies (Flyaa) was utilised, (Ma et al., 2020; Piper et al., 2017). Finally, for practical reasons we used cholesterol in the diet as opposed to ergosterol, because it is easily accessible, and where studied, has been shown to be adequate to support drosophila adult nutrition to the same extend as a yeast-based diet (Piper *et al.*, 2014).

**Figure 1.**
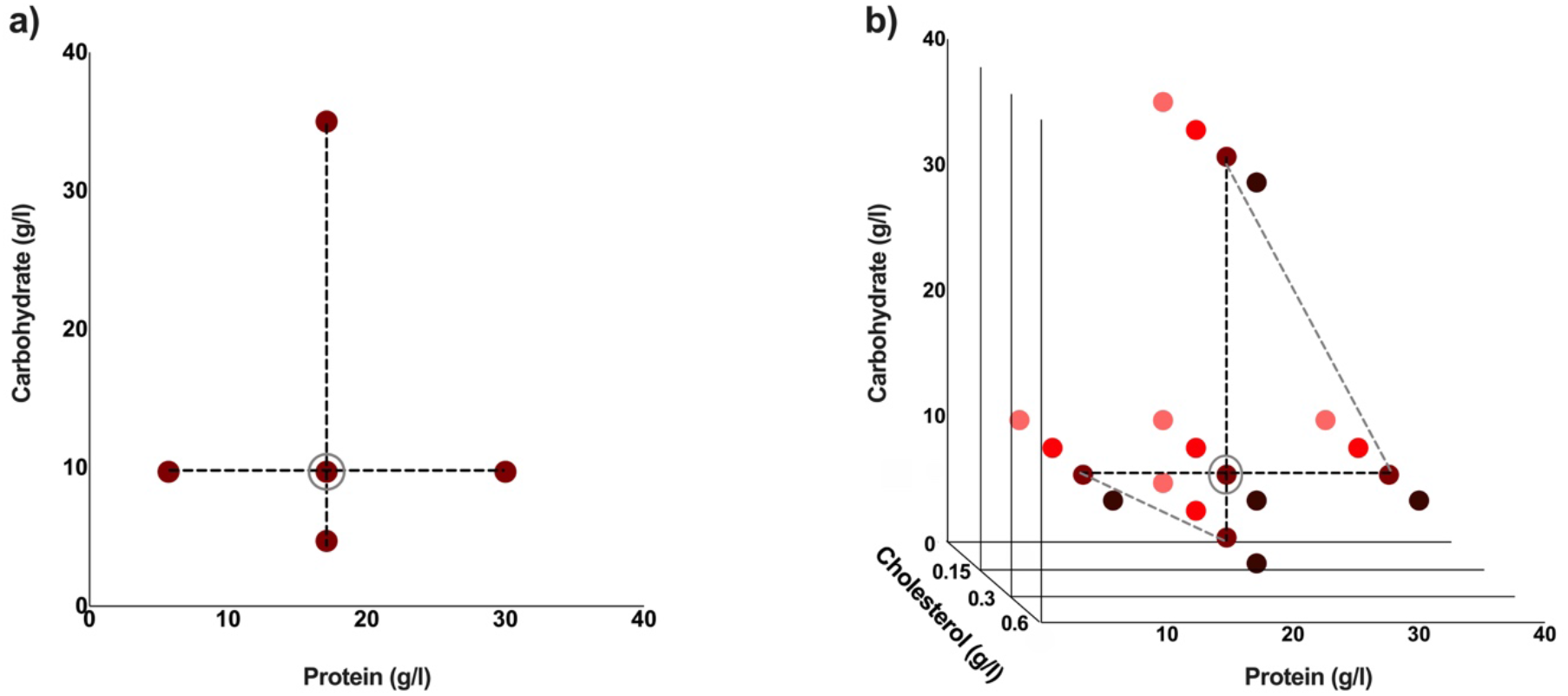
Experimental diets used are indicated by coloured dots. These diets have varying protein : carbohydrate ratios. This makes a total of five different experimental diets (a). The standard cholesterol concentration is 0.3g/l. Three additional cholesterol concentrations were used for each of the five protein : carbohydrate ratios to make a total of 20 different experimental diets (b). Diets which are either protein constant or carbohydrate constant are connected by black dotted lines, and diets with comparable caloric concentrations are connected by grey dotted lines (a, b). The standard diet used in our laboratory is circled in grey (a, b).

**Table 1.** Protein: carbohydrate ratio, along with the nutrient densities, cholesterol concentration and caloric content, for all synthetic experimental diets used. In the holidic media, amino acids are used to make up protein equivalents. To convert amino acids to protein equivalents, we used the molar quantities of nitrogen and the assumption that N makes up 16% of whole proteins (Imafidon & Sosulski, 1990). Calories were calculated using the method described in Southgate and Durnin (1970).

| Diet | Protein :<br>carbohydrate<br>equivalent | Sum<br>mass of<br>amino<br>acids (g/<br>L) | Equivalent<br>protein (g/L) | Carbohydrate (g/<br>L)<sup>2</sup> | Cholesterol<br>(g/l) | Estimated<br>caloric<br>content<br>(kcal/L) |
| --- | --- | --- | --- | --- | --- | --- |
| 1 | 1:0.6 | 5.25 | 4.7 | 17.1 | 0 | 90.68 |
| 2 | 1:0.6 | 5.25 | 4.7 | 17.1 | 0.15 | 90.68 |
| 3 | 1:0.6 | 5.25 | 4.7 | 17.1 | 0.3 | 90.68 |
| 4 | 1:0.6 | 5.25 | 4.7 | 17.1 | 0.6 | 90.68 |
| 5 | 1:0.6 | 10.74 | 9.7 | 35 | 0 | 186.05 |
| 6 | 1:0.6 | 10.74 | 9.7 | 35 | 0.15 | 186.05 |
| 7 | 1:0.6 | 10.74 | 9.7 | 35 | 0.3 | 186.05 |
| 8 | 1:0.6 | 10.74 | 9.7 | 35 | 0.6 | 186.05 |
| 9 | 1:1.8 | 10.74 | 9.7 | 17.1 | 0 | 118.93 |
| 10 | 1:1.8 | 10.74 | 9.7 | 17.1 | 0.15 | 118.93 |
| 11<sup>1</sup> | 1:1.8 | 10.74 | 9.7 | 17.1 | 0.3 | 118.93 |
| 12 | 1:1.8 | 10.74 | 9.7 | 17.1 | 0.6 | 118.93 |
| 13 | 1:3.6 | 33.1 | 30 | 17.1 | 0 | 233.63 |
| 14 | 1:3.6 | 33.1 | 30 | 17.1 | 0.15 | 233.63 |
| 15 | 1:3.6 | 33.1 | 30 | 17.1 | 0.3 | 233.63 |
| 16 | 1:3.6 | 33.1 | 30 | 17.1 | 0.6 | 233.63 |
| 17 | 1:3.6 | 10.74 | 9.7 | 5.7 | 0 | 76.18 |
| 18 | 1:3.6 | 10.74 | 9.7 | 5.7 | 0.15 | 76.18 |
| 19 | 1:3.6 | 10.74 | 9.7 | 5.7 | 0.3 | 76.18 |
| 20 | 1:3.6 | 10.74 | 9.7 | 5.7 | 0.6 | 76.18 |

<sup>1</sup> Standard diet.
<sup>2</sup> Carbohydrate is added to the diet as sucrose

The same methods for making the holidic medium described above were used to make all diets used in the rapamycin experiment. In this case however 18.9g/l protein: 17.1g/l carbohydrate were used. Cholesterol was supplemented so that its final concentration in food was 0.1g/l and rapamycin was added to a final concentration in the diet of 10uM. Diets were either un-supplemented, supplemented with cholesterol, rapamycin, or both.

#### Yeast based experiments

Four sugar/yeast (SY) diets were created using sucrose (Bundaberg Sugar, Melbourne Distuributors) and either whole yeast autolysate (MP Biomedicals, LLC, #903312) or yeast extract (Bacto Yeast Extract, #212750). These diets correspond to previously published conditions for fully fed and dietary restriction (dietary restriction) conditions (Bass et al., 2007; Katewa et al., 2016; Mair et al., 2005). The fully fed diets contained, per litre, 50g sucrose and 100g autolysed yeast or 50g sucrose, 50g yeast extract plus 86g of cornmeal (The Full Pantry, Victoria, Australia). The dietary restriction diets contained, per litre 50g sucrose and 200g autolysed yeast or 50g sucrose, 5g yeast extract plus 86g cornmeal. To each of these diets, we added cholesterol (Glentham Life Sciences, GEO100, #100IEZ) at a concentration of either 0 or 0.3g/l (Figure 2, Table 3). Cholesterol was added to all diets as a powder which was mixed in with all other dry ingredients prior to cooking. This gave us a total of four experimental diets per yeast.

### Statistical analyses

All statistical analyses were performed using R (version 3.3.0, available from http://www.R-project.org/). Linear mixed effect models were used to analyse all data obtained using the holidic media, along with egg data for yeast based diets. For the analysis of data obtained using the holidic media, a model reduction was performed by stepwise removal of the most complex non-significant term until any further removal significantly reduced the model fit. Log rank tests were used to compare the survival curves in the rapamycin experiment and yeast based dietary experiments. Finally, an ANOVA was used to analyse egg laying results for the rapamycin experiment. Plots were made in Graphpad Prism (version 8.4.2)

## Acknowledgements

We would like to thank Amy Dedman from Monash University for technical assistance, Xiaoli He and Dr. Mingyao Yang (University College London at the time) for contributions to the early phase of these experiments, as well as Lisa Rapley (Monash University) for help with eggs counts.

