## Supplementary Table 1 for "A dietary sterol trade off determines lifespan responses to dietary restriction in *Drosophila melanogaster*"

**Supplementary Table 1.** The relative proportions of each amino acid in the FLYaa amino acid mixture used in this study.

| Amino acids | | FLYaa |
|---|---|---|
| Essential amino acids | Phenylanine | 0.037 |
| | Histidine | 0.026 |
| | Isoleucine | 0.052 |
| | Lysine | 0.057 |
| | Leucine | 0.094 |
| | Methionine | 0.025 |
| | Arginine | 0.057 |
| | Threonine | 0.056 |
| | Valine | 0.062 |
| | Tryptophan | 0.010 |
| Non essential amino acids | | |
| | Alanine | 0.075 |
| | Cysteine | 0.017 |
| | Aspartate | 0.053 |
| | Glutamate | 0.063 |
| | Glycine | 0.062 |
| | Asparagine | 0.047 |
| | Proline | 0.052 |
| | Glutamine | 0.046 |
| | Serine | 0.079 |
| | Tyrosine | 0.031 |
