## Supplementary Figure 1 for "A dietary sterol trade off determines lifespan responses to dietary restriction in *Drosophila melanogaster*"

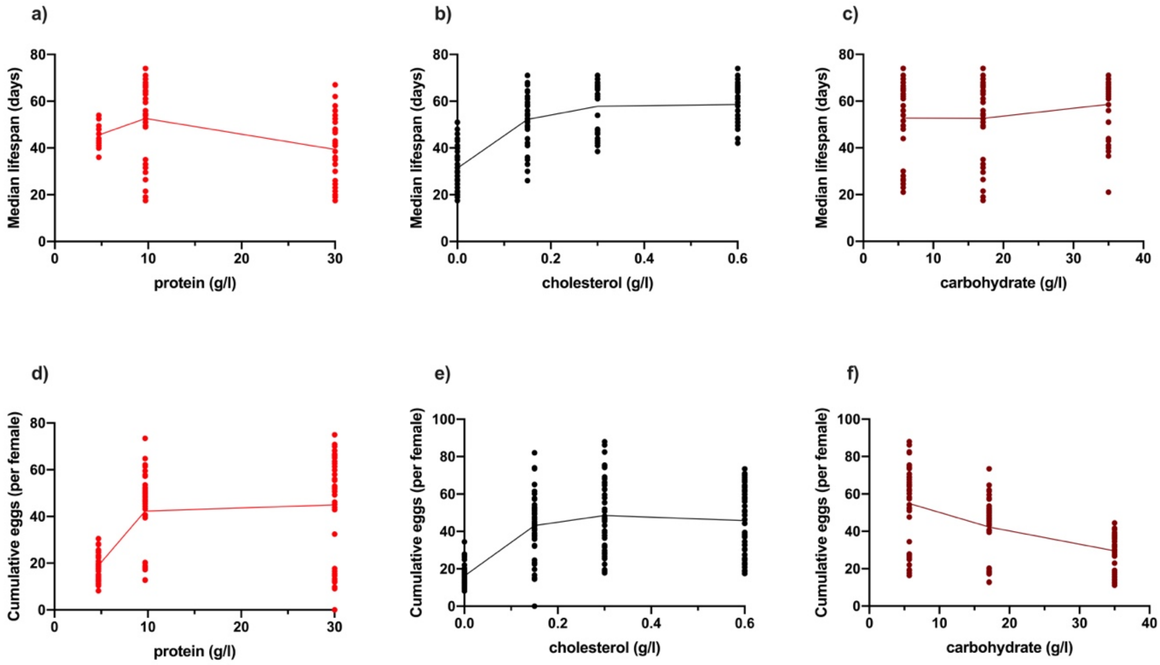


**Supplementary Figure 1.**

Changing dietary protein and cholesterol concentrations modify *Drosophila* lifespan (a-c), while changing protein, cholesterol and carbohydrate concentrations modify egg production (d-f). Diets varying in protein and carbohydrate concentration were each made at 4 levels of cholesterol (b , e) 0g/l, 0.15g/l, 0.3g/l and 0.6g/l. (a) Lifespan is reduced at both low and high protein concentrations, (b) was improved by increasing dietary cholesterol concentration, and (c) was unchanged by changing dietary carbohydrate concentration. Egg laying was improved by increasing protein (d) and cholesterol (e) concentrations, but decreased with carbohydrate concentration (f).
