## Supplementary Table 8 for "A dietary sterol trade off determines lifespan responses to dietary restriction in *Drosophila melanogaster*"

Effects on median lifespan (days) of cumulative eggs per female, cholesterol, cholesterol^2 and the interaction between cumulative egg production and cholesterol. Cumulative egg production and cholesterol had a significant positive effect on median lifespan, while cholesterol^2 had a significant negative effect on median lifespan. Data were analysed using a linear model with mixed effects, with vial as a random effect.

| **Variable** | **Estimate** | **Std. Error** | **t value** | **Pr (>Chisq)** |
|---|---|---|---|---|
| Cumulative eggs | 0.037 | 0.137 | 0.267 | 0.016 ** |
| Cholesterol | 115.620 | 27.262 | 4.241 | < 0.001 *** |
| Cholesterol$^2$ | -148.892 | 45.292 | -3.287 | < 0.001 *** |
| Cumulative eggs : Cholesterol | 0.358 | 0.898 | 0.398 | 0.691 |
| Cumulative eggs : Cholesterol$^2$ | 0.046 | 1.247 | -0.037 | 0.970 |
