## Supplementary Table 7 for "A dietary sterol trade off determines lifespan responses to dietary restriction in *Drosophila melanogaster*"

Effects on cumulative eggs per female of P:C ratio, cholesterol, cholesterol^2 and the interaction between P:C ratio and cholesterol. Each of the main effects had a significant positive effect on egg production, and the amount of cholesterol significantly modified how P:C affected egg laying. Data were analysed using a linear model with mixed effects, with vial as a random effect.

| **Variable** | **Estimate** | **Std. Error** | **t value** | **Pr (>Chisq)** |
|---|---|---|---|---|
| P:C | 2.226 | 2.348 | 0.948 | < 0.001 *** |
| Cholesterol | 69.478 | 22.638 | 3.069 | < 0.001 *** |
| Cholesterol$^2$ | -90.500 | 35.178 | -2.573 | < 0.001 *** |
| P:C : Cholesterol | 115.538 | 19.991 | 5.779 | < 0.001 *** |
| P:C : Cholesterol$^2$ | -135.520 | 31.095 | -4.358 | < 0.001 *** |
