## Supplementary Table 6 for "A dietary sterol trade off determines lifespan responses to dietary restriction in *Drosophila melanogaster*"

Minimum adequate model describing the effects of protein, protein^2 carbohydrate, carbohydrate^2, cholesterol, cholesterol^2 and, where appropriate, their interactive effects on cumulative eggs per female. Data were analysed using a linear model with mixed effects, with vial as a random effect.

| **Variable** | **Estimate** | **Std. Error** | **t value** | **Pr (>Chisq)** |
|---|---|---|---|---|
| Protein | 5.598 | 0.752 | 7.443 | <0.001*** |
| $Protein^2$ | -0.137 | 0.018 | -7.523 | <0.001*** |
| Carbohydrate | -0.253 | 0.129 | -1.956 | <0.001*** |
| Cholesterol | 218.625 | 27.209 | 8.035 | <0.001*** |
| $Cholesterol^2$ | -343.733 | 35.242 | -9.754 | <0.001*** |
| Protein : cholesterol | 6.239 | 2.189 | 2.850 | 0.004** |
| $Protein^2$: cholesterol | -0.104 | 0.053 | -1.950 | 0.051 |
| Carbohydrate : cholesterol | -5.427 | 1.106 | -4.905 | <0.001*** |
| Carbohydrate : $cholesterol^2$ | 7.000 | 1.722 | 4.066 | <0.001*** |
