## Supplementary Table 5 for "A dietary sterol trade off determines lifespan responses to dietary restriction in *Drosophila melanogaster*"

Minimum adequate model describing the effects of protein, protein^2 carbohydrate, carbohydrate^2, cholesterol, cholesterol^2 and, where appropriate, their interactive effects on median lifespan (days). Data were analysed using a linear model with mixed effects, with vial as a random effect.

| **Variable** | **Estimate** | **Std. Error** | **t value** | **Pr (>Chisq)** |
|---|---|---|---|---|
| Protein | -2.21 | 5.882 | -3.764 | <0.001*** |
| Protein$^2$ | 3.732 | 1.448 | 2.578 | <0.001*** |
| Carbohydrate | -2.365 | 2.592 | -0.912 | 0.362 |
| Carbohydrate$^2$ | 1.794 | 6.248 | 2.871 | 0.072 |
| Cholesterol | 1.989 | 1.298 | 1.533 | <0.001*** |
| Cholesterol$^2$ | -1.607 | 1.122 | -14.324 | <0.001*** |
| Protein: cholesterol | 1.801 | 1.560 | 11.545 | <0.001*** |
| Protein$^2$ : cholesterol | -4.135 | 3.800 | -10.882 | <0.001*** |
| Carbohydrate$^2$ : cholesterol | -2.763 | 5.154 | -5.362 | <0.001*** |
