## Supplementary Table 4 for "A dietary sterol trade off determines lifespan responses to dietary restriction in *Drosophila melanogaster*"

Estimates from a linear mixed effects model to explain the effects of protein and carbohydrate on lifespan (median lifespan (days)), with vial as a random effect. While variations in carbohydrate had no effect on lifespan, increasing doses of protein resulted in a significant change in lifespan. Visual inspection of the data agreed with out past experience with these diets (Piper et al, 2014 & 2017) that the lifespan response was best modelled by the quadratic term for protein ($Protein^2$) since lifespan peaked at our intermediate protein dose and fell away at both higher and lower doses. The quadratic term for Carbohydrate was thus also added to maintain balance in the model ($Carbohydrate^2$). The interaction between protein and carbohydrate is not included in any of our analyses because these terms were not co-varied in a balanced way in our experimental design.

| **Variable** | **Estimate** | **Std. Error** | **t value** | **Pr (>Chisq)** |
|---|---|---|---|---|
| Protein | 7.088 | 0.642 | 11.6056 | < 0.001*** |
| $Protein^2$ | -0.180 | 0.015 | -11.824 | < 0.001*** |
| Carbohydrate | -0.032 | 0.373 | -0.085 | 0.932 |
| $Carbohydrate^2$ | 0.001 | 0.009 | 0.083 | 0.934 |
