## Supplementary Table 3 for "A dietary sterol trade off determines lifespan responses to dietary restriction in *Drosophila melanogaster*"

Estimates from a linear mixed effects model to explain the effects of protein and carbohydrate on cumulative eggs laid per female, with vial as a random effect. Decreasing doses of carbohydrate and increasing doses of protein resulted in significantly increased egg production.

| **Variable** | **Estimate** | **Std. Error** | **t value** | **Pr (>Chisq)** |
|---|---|---|---|---|
| Protein | 0.809 | 0.204 | 3.964 | < 0.001*** |
| Carbohydrate | -1.315 | 0.2178 | -6.035 | < 0.001*** |
