## Supplementary Table 2 for "A dietary sterol trade off determines lifespan responses to dietary restriction in *Drosophila melanogaster*"

**Supplementary Table 2.** Protein: carbohydrate (P:C) ratio, along with the nutrient densities, cholesterol supplementation value and caloric content, for all yeast based experimental diets used.

| Diet | P:C equivalent | Sum mass of yeast (g/L) | Equivalent protein (g/L)[2] | Equivalent carbohydrate (g/ L)[3] | Cholesterol supplementation (g/l)[4] | Estimated caloric content (kcal/L) |
|---|---|---|---|---|---|---|
| 1[1] | 1:34.17 | 5 | 3.4 | 116.2 | 0 | 457 |
| 2[1] | 1:34.17 | 5 | 3.4 | 116.2 | 0.3 | 457 |
| 3[1] | 1:3.46 | 50 | 33.6 | 116.2 | 0 | 646 |
| 4[1] | 1:3.46 | 50 | 33.6 | 116.2 | 0.3 | 646 |
| 5 | 1:1.5 | 100 | 45 | 68 | 0 | 545 |
| 6 | 1:1.5 | 100 | 45 | 68 | 0.3 | 545 |
| 7 | 1:0.9 | 200 | 90 | 86 | 0 | 867 |
| 8 | 1:0.9 | 200 | 90 | 86 | 0.3 | 867 |

[1] Yeast extract media.

[2] Protein is added to the diet as autolysed yeast or yeast extract and cornmeal

[3] Carbohydrate is added to the diet as sucrose and cornmeal (in yeast extract media).

[4] Supplemented in addition to naturally occurring concentration.
